# Biologically realistic mean field model of spiking neural networks with fast and slow inhibitory synapses

**DOI:** 10.1101/2025.02.28.640853

**Authors:** Claudio Di Geronimo, Alain Destexhe, Matteo di Volo

## Abstract

We present a mean field model for a spiking neural network of excitatory and inhibitory neurons with fast GABA_**A**_ and nonlinear slow GABA_**B**_ inhibitory conductance-based synapses. This mean field model can predict the spontaneous and evoked response of the network to external stimulation in asynchronous irregular regimes. The model displays theta oscillations for sufficiently strong GABA_**B**_ conductance. Optogenetic activation of interneurons and an increase of GABA_**B**_ conductance caused opposite effects on the emergence of gamma oscillations in the model. In agreement with direct numerical simulations of neural networks and experimental data, the mean field model predicts that an increase of GABA_**B**_ conductance reduces gamma oscillations. Furthermore, the slow dynamics of GABA_**B**_ synapses regulates the appearance and duration of transient gamma oscillations, namely gamma bursts, in the mean field model. Finally, we show that nonlinear GABA_**B**_ synapses play a major role to stabilize the network from the emergence of epileptic seizures.

## 1 Introduction

Neural networks in the cerebral cortex exhibit a wide range of dynamical regimes, from asynchronous dynamics to synchronous irregular regimes characterized by oscillations at the scale of a population of neurons [1, 2]. Oscillations at the population scale, detected through mesoscopic signals such as Local Field Potentials (LFPs), are typically characterized by irregular and low firing activity of neurons at the microscopic scale [3] and have been shown to play an important role in brain computation. In particular, gamma oscillations (40-120 Hz) have been found to be implicated in cognitive functions, more specifically in attention [4, 5]. At the same time, slower oscillations such as theta oscillations (4-12 Hz) [6] have been shown to coordinate attention selection and memory [7, 8]. Moreover, the cross-frequency coupling between these different rhythms controls the functional connectivity and the large scale interactions in the brain [9]. Nevertheless, it is still unclear today what are the circuit mechanisms behind the different oscillations observed in cortical networks’ dynamics.

Low dimensional mean field models are powerful tools to propose potential circuit mechanisms regulating neural networks’ dynamics and stability. Characterized by few differential equations for population variables, mean field models define a formal link between the microscopic structure and the mesoscopic emergent dynamics of neural networks. Moreover, being low dimensional, mean field models also serve as the building block to model large scale dynamics at the whole brain scale [10]. Since the pioneer work by Wilson and Cowan [11] there has been intense research in the area of mean field models applied to the cortical dynamics [12–16]. While the first mean field models were purely phenomenological, many efforts have been made to develop mean field models that actually represent the population dynamics of a spiking neural network. This propriety is important because it allows us to validate mean field model’s predictions. Nevertheless, given the complexity and nonlinearities of networks of spiking neurons, such dimensional reduction procedure is mathematically challenging and mean field models need to rely on assumptions on the microscopic structure of the network. These assumptions often limit the inclusion of biologically realistic ingredients such as network’s structure, neurons diversity or conductance-based interactions. Recently, an exact mean field model has been introduced for Quadratic integrate and fire neurons [17]. This approach has been shown to be able to model gamma oscillations and theta-gamma coupling [18], including the bursting features of gamma oscillations [19]. Nevertheless, this approach is limited to QIF neurons, all-to-all networks, and Cauchy distributions of heterogeneities. Recent works have shown that new dynamical regimes emerge instead in sparse networks [20, 21] or in the presence of non-Cauchy distributions [22].

In this paper we consider instead networks with biologically realistic ingredients, such as sparse network connections, conductance-based interactions and different spike shape between regular spiking excitatory and fast spiking inhibitory neurons. We refer to our model as biologically realistic, because it integrates these ingredients. In order to derive a mean field model of this network we employ a different approach, based on master equation methodology [14] that has been shown to be extendable to different neurons types (from Hodgkin Huxley to Morris-Lecar) [23], to account for any distribution of heterogeneity [24] and to predict the population dynamics of a sparse network of spiking neurons [25]. This approach has been successfully employed for modeling asynchronous dynamics, slow oscillations [24] and recently gamma oscillations with the addition of synaptic delay [26]. Nevertheless, these previous mean field models consider only two types of synapses, fast AMPA and *GABA*_*A*_ receptors.

In this work we are interested in the role of slow and nonlinear inhibitory synapses such as *GABA*_*B*_ receptors. We thus developed a new mean field model including the effect of nonlinear slow *GABA*_*B*_ receptors. We have employed an adiabatic approximation, already tested effective in this model for spike frequency adaptation [25], and we have performed mean field approximation to account for the experimentally observed nonlinearity of *GABA*_*B*_ receptors. This method allows to account for non-linear slow inhibitory synapses. We show that this mean field model can reproduce the network response to external stimulation and that it predicts the emergence of theta oscillations at 4-5 Hz appearing for sufficiently strong *GABA*_*B*_ conductance. Interestingly, this new mean-field model predicts distinct effects between the direct stimulation of inhibitory neurons and the enhancement of *GABA*_*B*_ receptors in the emergence of gamma oscillations. Indeed, while the stimulation of inhibitory neurons does increases the power of gamma oscillations, an increase of *GABA*_*B*_ conductance eliminates gamma oscillations. This results shows that increasing inhibition can have different effects, depending on how inhibition is increased in a network of neurons. These predictions are consistent with direct network simulations of spiking neural networks and with experimental observations. Indeed, *GABA*_*B*_ agonist baclofen has been shown to suppress gamma oscillations in hippocampal slices [27]. By employing electrical stimulation, it is in fact found that synaptically released GABA was capable of modulating gamma frequency oscillations via activation of *GABA*_*B*_ receptors [27]. On the contrary, optogenetic stimulation of parvalbumin-positive (PV) interneurons has been shown to increase the power of gamma oscillations [28, 29]. We also employ our mean field model to model gamma bursts, i.e. periods of few gamma cycles separated by asynchronous activity [30]. We show, through our mean field model, that *GABA*_*B*_ dynamics is strongly linked with gamma oscillatory dynamics. In fact, the appearance and the duration of bursts of gamma oscillations are driven by the temporal evolution and the decay of *GABA*_*B*_ receptors. Finally, we show that the nonlinearity of *GABA*_*B*_ synapses (activating only when a certain amount of presynaptic spikes arrive in synchrony [31]) allows the network to stabilize and avoid epileptic seizures.

In summary, this mean field model is a good candidate to model different dynamical regimes in mesoscopic signals, from asynchronous dynamics to slow and fast oscillations. Importantly, it can be used to test pharmacological action on different inhibitory receptors at the mesoscopic scale and, when integrated in large scale models of the whole brain, it can produce valuable predictions on the effect of such manipulation for large scale activity in the brain and epileptic spreading.

## 2 Materials and Methods

### 2.1 Spiking neural network model

We consider a network of N=10000 Adaptive Exponential Integrate and Fire neurons. *N*_*e*_ = 8000 are excitatory Regular Spiking (RS) neurons and the remaining *N*_*i*_ = 2000 3 are inhibitory Fast Spiking (FS). Neurons are randomly connected over a sparse network with a probability of connection between two neurons of *p* = 5%. The temporal evolution of neurons’ membrane potential is described by the following differential equations:

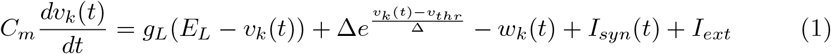

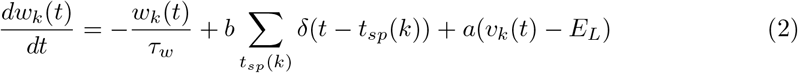

At spike time (v_k_ *>* v_thr_) : *v*_*k*_ →− *v*_*rest*_, *w*_*k*_ →− *w*_*k*_ + *b*,

where *C*_*m*_ = 200pF is the membrane capacity, *v*_*k*_ is the voltage of neuron *k*, and whenever *v*_*k*_ *> v*_*thr*_ = −30mV, *v*_*k*_ is reset to the resetting voltage *v*_*rest*_ = −65mV and fixed to that value for a refractory time of *T*_*refr*_ = 5ms. The variable *w*_*k*_ is a self generated current that simulates the adaptation of a neuron after a spike. Due to physiological insight only excitatory neurons have adaptation with values of *a* = 0, *b* = 20pA, *τ*_*w*_ = 500ms (*a* = *b* = 0 for all inhibitory neurons). The leak conductance used is *g*_*l*_ = 10nS and *E*_*l*_ = − 65mV is the leakage reversal potential. The exponential term is slightly different for RS (Δ = 2mV) and FS neurons (Δ = 0.5mV) while the effective threshold is the same for all neurons: *v*_*thr*_ = − 50mV. *I*_*ext*_ describes an external current that is set to zero if not stated otherwise. We will employ this in Sec. 3.5 to manually activate inhibitory neurons as in optogenetic experiments. *I*_*syn*_ describes all types of presynaptic inputs. In particular, in conductance-based models, such a current is generated by changes in conductances when presynaptic neurons fire:

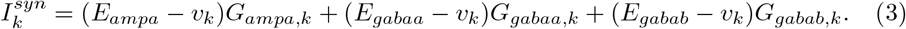

In our model we will consider 3 types of receptors that can change the conductances of the membrane with the opening of associated ionic channels. *G*_*ampa*_ is increased by incoming excitatory spikes, while *G*_*gabaa*_ and *G*_*gabab*_ are increased by spikes of presynaptic inhibitory neurons. Each channel has its own reversal potential: *E*_*ampa*_ = 0mV, *E*_*gabaa*_ = −80mV and *E*_*gabab*_ = −90mV. The time evolution of individual conductances is modeled as a decaying exponential function increased of a fixed amount *Q*_*x*_ at each presynaptic spike:

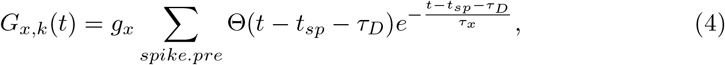

with *x* = *ampa, gabaa* or *gabab*, Θ is the Heaviside function and *τ*_*D*_ the synaptic delay. In this work we consider either instantaneous synapses (Section 3.1-3.4), i.e. with a delay of synaptic transmission *τ*_*D*_ set to 0ms, either homogeneous delay *τ*_*D*_ = 3ms (Section 3.5-3.6). It is known that *GABA*_*B*_ receptor have non linear activation. Indeed, receptors are fully open only when a sufficient number of presynaptic spikes occur within short time intervals, typically 4-10 spikes in 100ms [31]. In order to include such nonlinearity, we consider that the conductance *g*_*gabab*_ depends on the amount of inhibitory inputs received in the last 100ms, that we approximate as 0.1*K*_*i*_*ν*_*i*_, where *K*_*i*_ = *pN*_*I*_ is the average number of inhibitory presynaptic connections and *ν*_*i*_ (*ν*_*e*_) the inhibitory (excitatory) population firing rate. More specifically, following [31] we consider the fraction of open receptors 𝒢 as

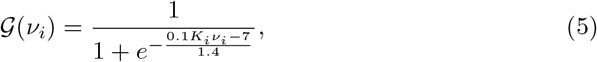

and the conductance of *GABA*_*B*_ synapses is thus *g*_*gabab*_ ·𝒢 (*ν*_*i*_), instead of simply *g*_*gabab*_. We determine the population activity of one cell type as

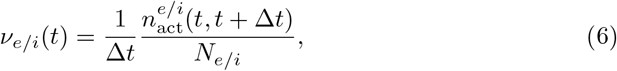

where *N*_*e/i*_ represents the number of neurons within the population. The term 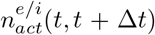 signifies the cumulative spike count across all neurons in the group within the time window (*t, t* + Δ*t*). In our simulations we use Δ*t* = 10ms. Synaptic parameters here employed are the same as in [25] (apart from those of *GABA*_*B*_ synapses). The *GABA*_*B*_ conductance *g*_*gabab*_ varies in our simulations. The constants of decay for the conductances are *τ*_*ampa*_ = *τ*_*gabaa*_ = 5ms and *τ*_*gabab*_ = 180ms if not stated otherwise. The quantal conductances are *g*_*ampa*_ = 1nS for RS to FS coupling, *g*_*ampa*_ = 1.05nS for RS to RS coupling and *g*_*gabaa*_ = 5nS for both FS to RS and FS to FS coupling. Each neuron receives an external drive, modeled by considering *K*_*e*_ = 400 RS neurons generating action potentials with a Poissonian statistic at a fixed frequency rate *ν*_*ext*_ = 2.5Hz. In Section 3.3 we have instead employed a time varying *ν*_*ext*_. First we used a rising and decaying input of the form:

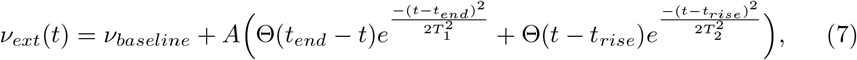

with *ν*_*baseline*_ = 2.5Hz, *T*_1_ = *T*_2_ = 20ms and *A* = 6Hz. Then we have also employed a simple sinusoidal wave form for *ν*_*ext*_ (see Sec. 3.3).

### 2.2 Mean Field Model

In order to derive the mean field model for this spiking neural network we adopt the master equation formalism first introduced in [32], based on a markovian assumption such that the network dynamics is memoryless over a time scale *T*. Nevertheless, as shown in [25] this assumption can be employed also in spiking neural networks where there are slower mean field processes such as spike frequency adaptation, by employing an adiabatic approximation. In our network we have two relatively slow processes: spike frequency adaptation and synaptic *GABA*_*B*_ conductances. We thus employ an adiabatic approximation for both processes. The equation for the population average *GABA*_*B*_ conductance, namely *G*_*b*_, is obtained by taking the derivative of Eq. (4) and then by applying the population average. All together, the first order equations for this mean field model are the following:

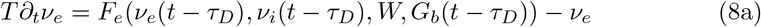

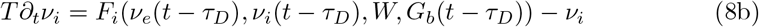

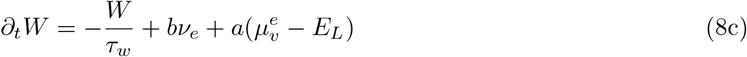

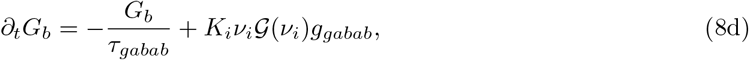

where *K*_*i*_ is the average number of presynaptic FS neurons, *F*_*e*,*i*_ are respectively the stationary transfer functions of RS and FS neurons. They express the output firing rate as a function of the rates of incoming excitatory and inhibitory neurons, the level of adaptation *W* and the level of *GABA*_*B*_ conductance *G*_*b*_. The impact of synaptic delay has been introduced following [26]. In the next chapter we will describe how to evaluate these transfer functions *F*_*e/i*_, their dependence on *GABA*_*B*_ conductance *G*_*b*_ and the mean population voltage 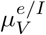 of excitatory/inhibitory neurons using a semianalytical derivation. In the following analysis we will use *T* = 10*ms* if not stated otherwise. We will often refer to stationary version of this mean field (i.e. *stationary mean field*), which is the version where we do not consider the temporal evolution of *GABA*_*B*_ synapses and *G*_*b*_(*t*) = *τ*_*gabab*_*K*_*i*_*g*_*gabab*_*ν*_*i*_(*t*), just like we do for fast *GABA*_*A*_ synapses.

### 2.3 Neurons transfer function (TF)

The theoretical analysis of the TF of RS and FS neurons is based on the shot-noise theory in [33], adapted to RS and FS neurons in [34, 35]. The assumption behind this approach is that it is possible to write the output firing rate of a neuron as a function of the subthreshold voltage statistic. We will estimate the average subthreshold voltage *µ*_*v*_, the standard deviation *σ*_*v*_ and the time correlation decay *τ*_*v*_ of neurons’ membrane potential. First we will report how to evaluate these quantities, and then how to relate them to the firing rate.

#### 2.3.1 From Input Rates to Subthreshold Voltage Moments

The mean membrane potential is obtained following the Methods described in [33], by estimating the stationary conductance given by an average synaptic Poissonian input rate (*ν*_*e*_, *ν*_*i*_):

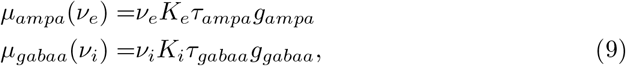

where *K*_*µ*_ = *pN*_*µ*_ represents the number of incoming synapses from excitatory (*µ* = *e*) or inhibitory (*µ* = *i*) neurons. These stationary conductances, together with the 6 instantaneous and time-varying *GABA*_*B*_ conductance *G*_*b*_, will define the effective time constant 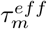 of the membrane:

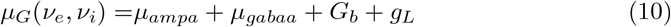

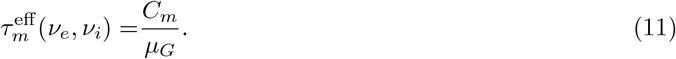

Given a specific value of *W* and *G*_*b*_, we can estimate the mean membrane potential as:

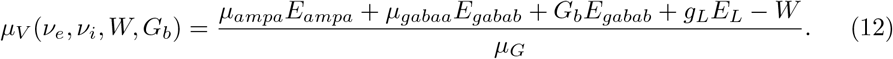

The calculation of *σ*_*v*_ and of *τ*_*v*_ is based on the assumption of Poissonian spiking statistics [35]. Each conductance gives an additional term to *σ*_*v*_ and *τ*_*v*_, so the effect of *GABA*_*B*_ synapses can be taken into account:

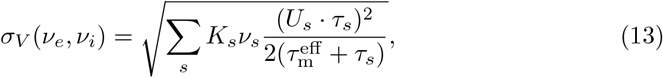

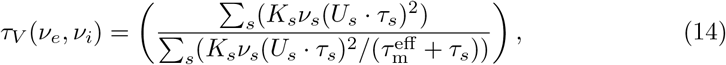

where *s* = *ampa, gabaa, gabab* and 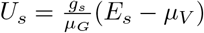.

#### 2.3.2 From Subthreshold Voltage Moments to the Output Firing Rate

We will now estimate the output firing rate of a neuron as a function of (*µ*_*V*_, *σ*_*V*_, *τ*_*V*_), using the following formula:

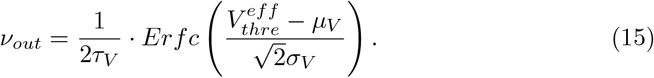

This formula is based on the hypothesis that the membrane potential follows a Gaussian distribution evolving over a timescale of *τ*_*V*_. We thus define the firing rate as the area of such distribution beyond the threshold 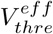. In this way we obtain an analytical framework to predict the firing rates as a function of (*µ*_*V*_, *σ*_*V*_, *τ*_*V*_). We then fit the function 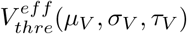 to single neurons’ simulations. This procedure overcomes the problem of finding a theoretical value for the threshold, which is not possible due to the nonlinear terms in the membrane potential equation, such as the exponential term in the membrane evolution and the conductance based input. The phenomenological threshold was taken as a second-order polynomial in the following way:

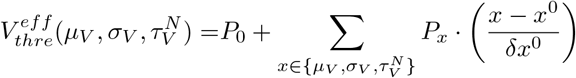

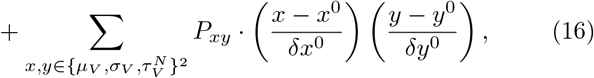

where we used the dimensionless timescale 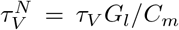. The fit of 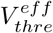 is evaluated using a simulation of single neuron with a Poissonian external input fixed and correcting the predicted output until it match the single neuron simulation. We used the following values to expand 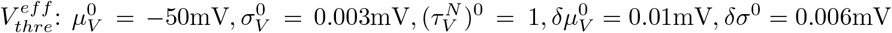 and *δ*(*τ* ^*N*^)^0^ = 0.2. In order to evaluate the values of P, we performed fitting using single neuron simulations with different values of *g*_*gabab*_. Results of the fit are reported in Table 1.

**Table 1.** Fit Parameters (Expressed in mV)

| Cell Type | $P_0$ | $P_{\mu_V}$ | $P_{\sigma_V}$ | $P_{\tau_V^N}$ | $P_{\mu_V^2}$ | $P_{\sigma_V^2}$ | $P_{(\tau_V^N)^2}$ | $P_{\mu_V \sigma_V}$ | $P_{\mu_V \tau_V^N}$ | $P_{\sigma_V \tau_V^N}$ |
| --- | --- | --- | --- | --- | --- | --- | --- | --- | --- | --- |
| FS | -49.47 | 0.01 | 7.81 | 0.66 | 8.90 | -0.53 | 1.43 | 0.85 | 13.68 | 0.25 |
| RS | -47.04 | 1.66 | -4.81 | 1.08 | 7.59 | 0.10 | -0.63 | 1.88 | 24.66 | 0.40 |

## 3 Results

### 3.1 Transfer function estimation

The building block of the mean field model in Eq.s (8) is the transfer function *F*. The transfer function represents the output firing rate of one RS or FS neuron when receiving excitatory and inhibitory Poissonian spikes at a rate *ν*_*e*_ and *ν*_*i*_. The transfer function is thus a propriety of one neuron type, not of a recurrent neural network (that we will treat in next section). In the formalism developed here the transfer function *F* depends on the rate of incoming excitatory and inhibitory Poissonian spikes *ν*_*e*_ and *ν*_*i*_, on the level of adaptation *W* and on the level of *GABA*_*B*_ conductance *G*_*b*_. In stationary conditions, those where the transfer function is estimated, adaptation and *GABA*_*B*_ conductance reach the value *W* ^∗^ = *bτ*_*w*_*ν*_*e*_ and 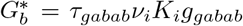, which depends only on *ν*_*e*_ and *ν*_*i*_. In Fig. 1a we show the transfer function for RS and FS neurons in absence of *GABA*_*B*_ synapses (*g*_*gabab*_ = 0nS). Dots stand for results of simulation of single neurons and the continuous line is the prediction based on the semi-analytical approach we employed to estimate *F*. We observe a very good agreement, as it was already proven in previous works [25, 34]. Notice that this semianalytical approach allows us to predict the output firing rate of a neuron for different parameters, without the need to perform again the fitting. The fitting is performed only once for one parameter setup, and thanks to the semi-analytic approach, we can theoretically predict isolated neurons firing rate for different parameters’ values. Indeed, even if the transfer function has been fitted for a specific parameter setup, we can test its capacity to predict neurons’ output activity in other parameters setup (i.e. for different values of *GABA*_*B*_ quantal conductance) synapses. This basically corresponds to testing the efficacy of our semi-analytic approach (see Section 2.3.1 of the Methods) in presence of time varying conductance based synapses. In Fig. 1b we report the transfer function for an increased level of *GABA*_*B*_ conductance, showing that the same analytical transfer function works also in presence of *GABA*_*B*_ synapses. We then extended this analysis by increasing systematically the amount of *GABA*_*B*_ conductance *g*_*gabab*_. In Fig. 1c we report the firing rate of RS and FS neurons for fixed incoming spike rates *ν*_*e*_ and *ν*_*i*_ but for different values of *GABA*_*B*_ quantal conductance *g*_*gabab*_.

**Fig. 1.**
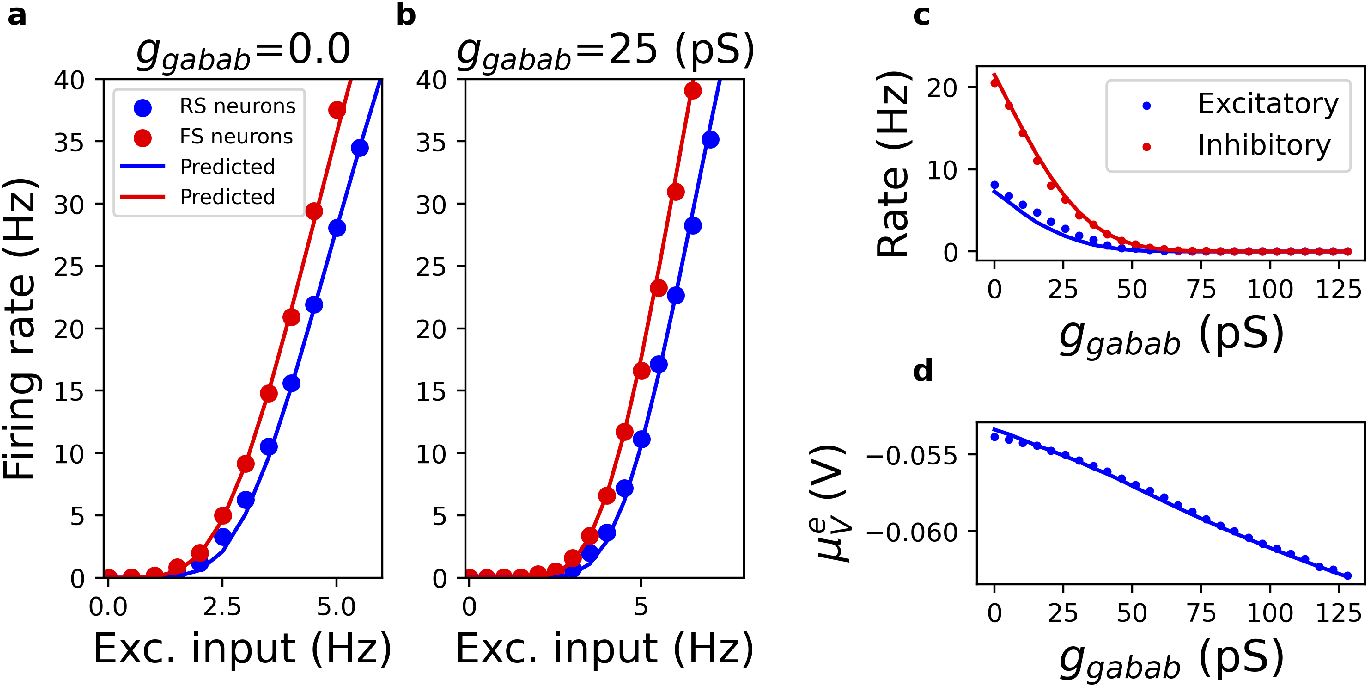
(a) Theoretical predictions (continuous lines) based on Eq. (15) compared with numerical simulations of the firing rates of RS (FS) neurons, represented as blue (red) dots, for different excitatory input rates *ν*_*e*_ and a fixed inhibitory input of *ν*_*i*_ = 8Hz. Quantal conductance of *GABA*_*B*_ synapses is *g*_*gabab*_ = 0pS. (b) Same as (a), but for *g*_*gabab*_ = 25pS. (c) RS (FS) neurons firing rates for fixed excitatory inputs *ν*_*e*_ = 4Hz and inhibitory input *ν*_*i*_ = 8Hz, shown as blue (red) dots for simulations and continuous line for theoretical predictions, as a function of *g*_*gabab*_. (d)Average membrane potential of RS neurons as function of *g*_*gabab*_ (same simulations as panel c).

As expected, the firing rate decreases increasing *g*_*gabab*_. Importantly, we observe a very good agreement between numerical simulations and theoretical predictions. Moreover, in Fig. 1d we observe that our theory (in particular Eq. 12, see Methods) allows us to predict the mean membrane potential of neurons for different level of *GABA*_*B*_ quantal conductance *g*_*gabab*_. We can thus now employ the transfer function to predict the emergent dynamics of recurrent neural networks of RS and FS neurons through the mean field Eq.s (8). Notice that even if we obtain a good theoretical transfer function for isolated neurons, this does not necessarily imply that we can predict the population dynamics of a sparsely connected network of interacting neurons. We show in the next chapters that the complete mean field Eq.s (8) here presented, are able to correctly predict sparse network dynamics.

### 3.2 Spontaneous activity in Asynchronous irregular regimes

Here, we study the emergent dynamics of a recurrently connected network composed of RS and FS neurons (see Methods section for details) in the absence of synaptic delay (*τ*_*D*_ = 0ms). In Fig. 2a we show the raster plot for RS neurons. Notice that we considered the presence of *GABA*_*B*_ synapses, with a quantal conductance *g*_*gabab*_ = 25pS. We observe an asynchronous irregular regime characterized by neurons with Poissonian firing statistics (we have verified that the coefficient of variation *CV* of neurons firing statistics is around *CV* = 1 for both RS and FS neurons, as expected for Poisson processes). Since the network dynamics is asynchronous, it should be represented by a fixed point in the mean-field model of Eq.s (8).

**Fig. 2.**
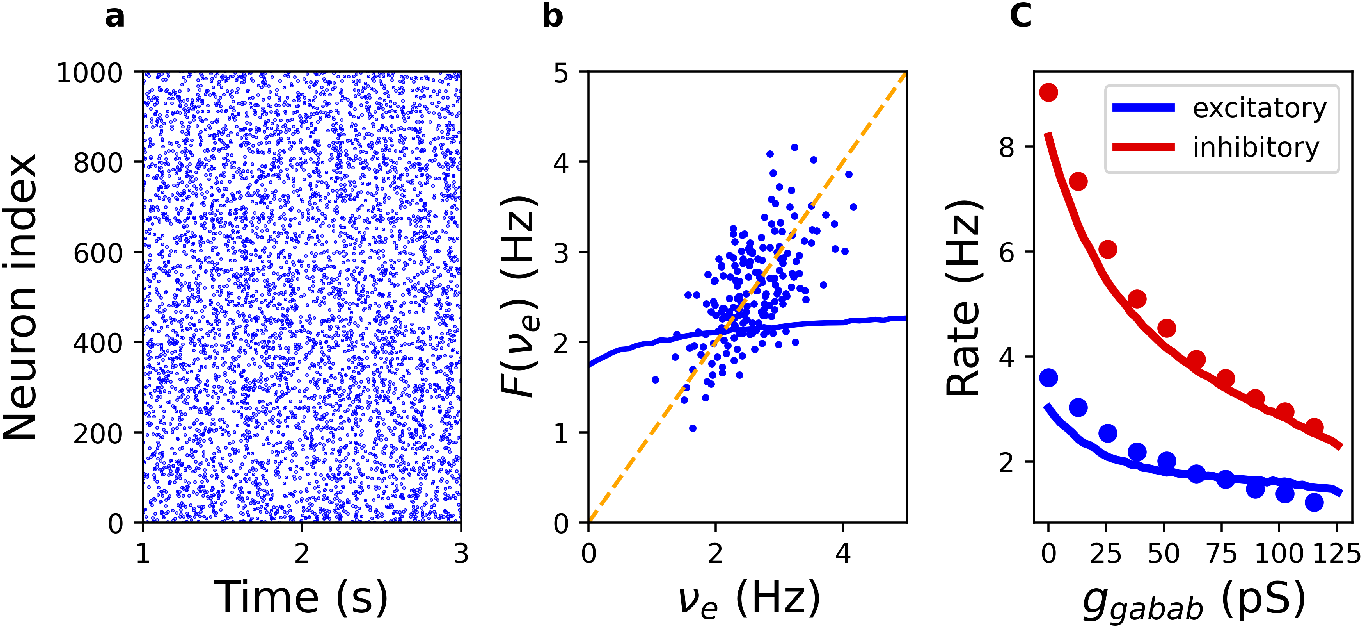
(a) Raster plot of excitatory neurons spiking activity for *g*_*gabab*_ = 25*pS*. The network simulation consists of 10,000 neurons, but we report only a subsample of 1,000 excitatory neurons. (b) Function *ℱ* (*ν*_*e*_) derived from the mean field equations (8) (see main text) is reported as a continuous blue line. The bisector is the orange line, while the dots represent the corresponding excitatory neurons firing rate in numerical simulations of the spiking neural network. (c) Comparison between the fixed point predicted by the mean field equations (continuous lines) and the values obtained from numerical simulations (dots), for different values of *g*_*gabab*_.

We thus report in Fig. 2b the consistency map for the theoretical excitatory neurons firing rate *ν*_*e*_ (see the continuous blue line). This map is obtained by estimating, for each value of excitatory input *ν*_*e*_, the corresponding stationary value of inhibitory neurons activity 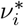, the adaptation variable *W* ^∗^, the *GABA*_*B*_ conductance 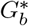 and corresponding excitatory firing rate 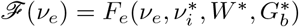. A consistent solution (fixed point) is obtained when *ℱ* (*ν*_*e*_) = *ν*_*e*_, i.e. when the blue line crosses the bisector (see the orange dashed line in Fig. 2a). As shown in Fig. 2b the mean field model predicts a stable fixed point, with a value of ~ 2 Hz. When reporting the values of *ν*_*e*_ at different time steps from numerical simulation of the neural network (see blue dots in Fig. 2b) we observe values with a mean of ~ 2.3 Hz characterized by natural fluctuations over time. These fluctuations are normally distributed [25] and we have verified that they vanish in the limit of infinite number of neurons. Moreover, the value of the mean rate in network smulations does not change for sufficiently large networks (i.e. more than 1,000 neurons). From these results we conclude that the mean field predictions are consistent with numerical simulations in terms of the mean activity of population rates. Notice that the mean formalism employed here can be extended to include equations predicting the dynamics of second order quantities, e.g. the amount of fluctuations in finite size networks [14, 25]. We limit here to discuss the prediction of the mean field in terms of the mean activity of population rates. We then asked if the prediction of the mean field model for stationary firing activity of excitatory and inhibitory neurons are consistent with network simulations increasing the strength of *GABA*_*B*_ conductances *g*_*gabab*_. In Fig. 2c we observe a very good agreement for both excitatory and inhibitory activity over a wide range of values of *g*_*gabab*_. In summary the mean field model correctly predicts stationary population activity for different values of *GABA*_*B*_ synaptic conductances.

### 3.3 Evoked activity in Asynchronous irregular regimes

We then investigated whether the mean field model correctly predicts network response to transient external stimulation. We thus introduced a time varying excitatory input frequency *ν*_*ext*_ and observed the response of the network.

In Fig. 3a and b we report the raster plot of excitatory neurons and the population firing rate *ν*_*e*_(*t*) in response to a rising stimuli at time *t*_*rise*_ = 3s, that goes back to the baseline value at time *t*_*end*_ = 5s, as described by Eq. (7) (see Methods section and the dashed line in the inset of Fig. 3b). Notice that we employ a network with excitatory and inhibitory neurons as described in the method section. We observe a very good agreement between the complete mean field model of Eq.s (8) (that includes the dynamics of *GABA*_*B*_ synapses) and the network simulation. Notice that before stimulation at *t*_*rise*_ = 3s the mean field predicts a rate ~ 2Hz and network simulation show a rate of ~ 2.3Hz with intrinsic fluctuations of ± 0.5Hz. The fluctuations observed in network simulations are due to the finite effects in our sparse network. We also report the results from a stationary version of this mean field model (dashed line). This stationary version is obtained by considering instantaneous *GABA*_*B*_ synapses, i.e. without considering their slow decay in time but only their tonic stationary value,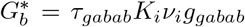. This is the same approach we employ for fast *GABA*_*A*_ and *AMPA* receptors. In the complete mean field instead, *G*_*b*_ follows its own differential equation, see Eq. (8d). Notice that when the population dynamics is stationary (i.e. non-evolving in time), the stationary mean field predicts the same fixed values for population rates to the complete mean field. In fact, before the stimulus onset at t=3s, the stationary mean field (dashed line) and the complete mean field (continuous line) are superimposed, and they predict very well the network activity. We observe instead that the complete mean field captures much better the transient response at stimulus onset, while the stationary mean field underestimates the response to the stimulus arrival at *t* = 3s and overestimates network activity at stimulus offset at time *t* = 5s. This is a direct consequence of the time scale of *GABA*_*B*_ synapses, that is taken into account in our mean field model. We have also tested a sinusoidal input at a frequency *f* = 2Hz. The results are shown in Fig.3c, where we observe that the complete mean field is capable to correctly reproduce the amplitude of the response entrained by the oscillatory input, at variance with the previous stationary mean field model. As we have shown in a previous work, the adiabatic approximation here proposed induced discrepancies between the mean field model predictions and the network simulation for too fast time scales [25]. This implies that whenever *τ*_*gabab*_ is too fast (typically faster that 100ms) and when the input frequency is too high (typically faster that 10 Hz), the mean field and the network simulation show a quantitative discrepancy due to the adiabatic approximation.

**Fig. 3.**
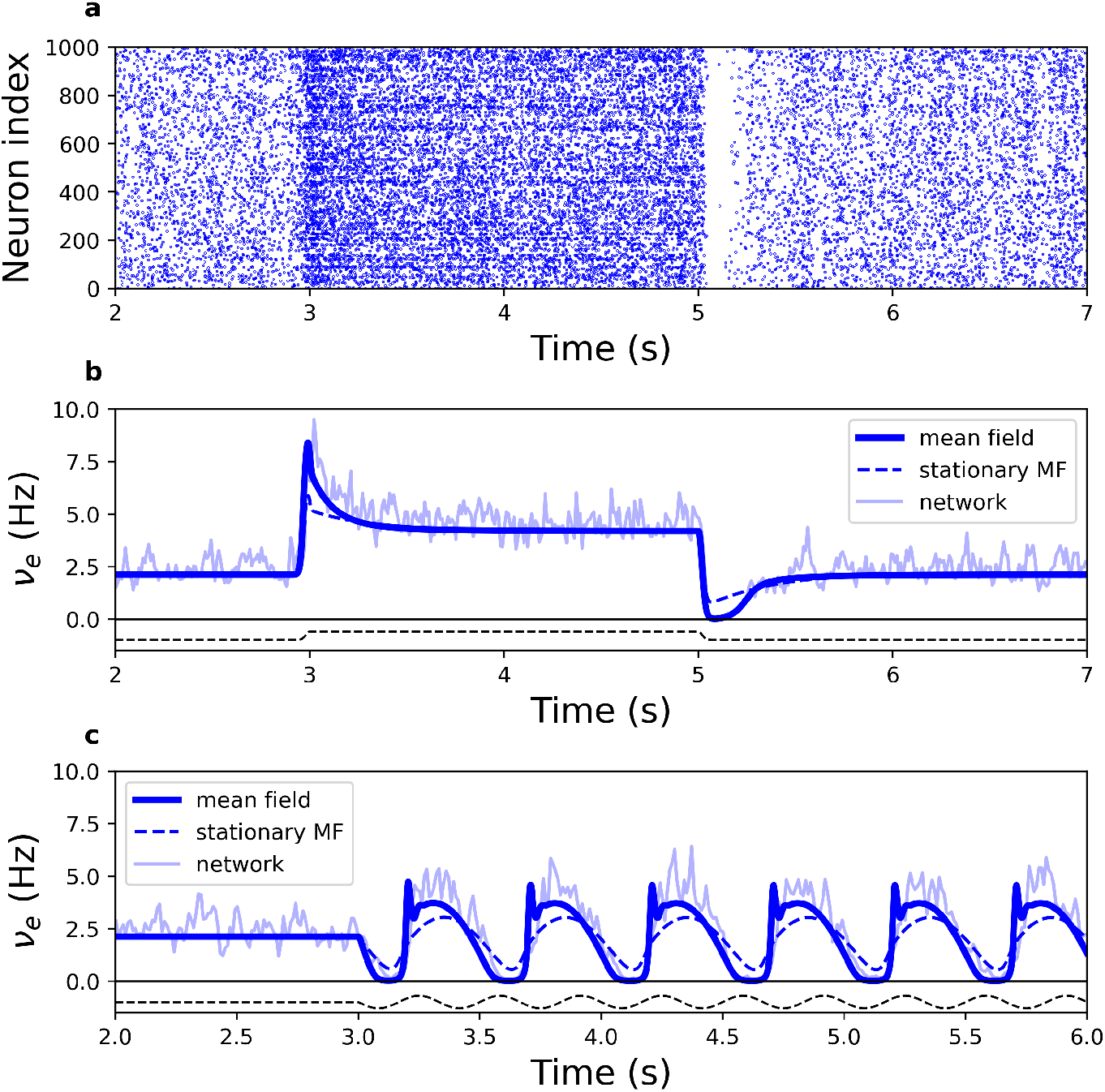
Comparison between mean field predictions, network simulations, and a stationary mean field where *G*_*b*_ is considered always equal to its stationary value. (a)Raster plot of excitatory neurons spiking activity for *g*_*gabab*_ = 25pS in response to an external input. The input is reported as a dashed line in panel b, and follows the Eq. (7), see Methods. The network simulation consists of 8,000 excitatory and 2,000 inhibitory neurons, we show here only a subsample of 1,000 excitatory neurons. (b) Corresponding RS neurons population rate *ν*_*e*_(*t*) (blue noisy line). The shape of the stimuli is represented by the black dotted line below the graph. The thick blue line is the mean field model prediction from Eq.s (8) while the dashed line is the stationary mean field model prediction (i.e. without *GABA*_*B*_ dynamics). (c) As (b) but for a sinusoidal input of frequency *f* = 2Hz and amplitude *A* = 1Hz (see Methods section).

### 3.4 Theta oscillations with GABA_B_ receptors

We have then employed the mean field model to predict if new dynamical regimes can be induced by the presence of *GABA*_*B*_ synapses. In Fig. 4a we report the standard deviation (over time) of the excitatory population firing rate, namely Σ(*ν*_*e*_). Red dots are obtained for the value of the external drive *ν*_*ext*_ = 2.5Hz used in Fig. 2 and

**Fig. 4.**
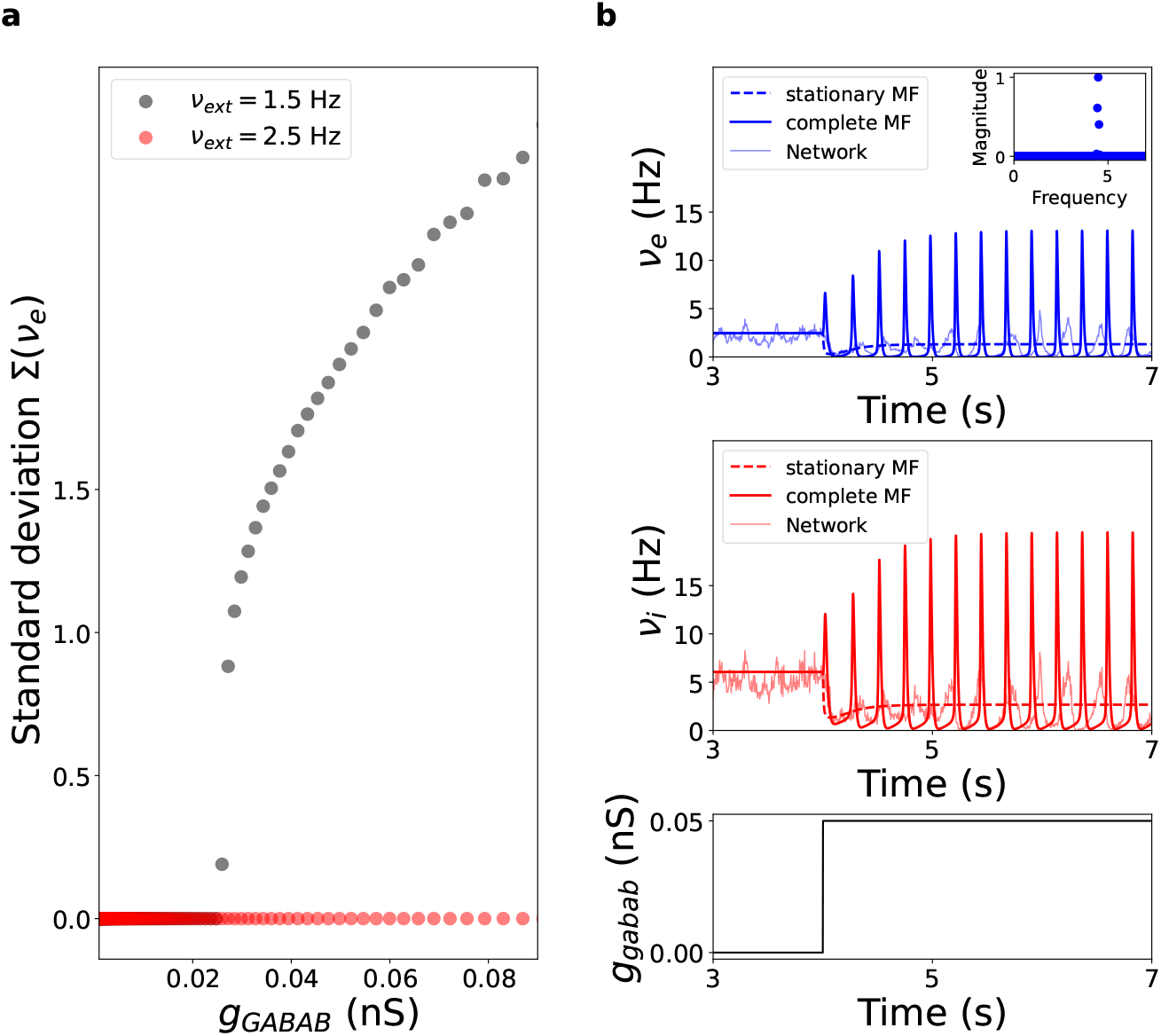
(a) The standard deviation (over time) Σ(*ν*_*e*_) of the excitatory neurons population rate *ν*_*e*_(*t*) estimated from the mean field model in Eq.s (8) as a function of *g*_*gabab*_ for *ν*_*ext*_ = 2.5 Hz (red dots) and *ν*_*ext*_ = 1.5 Hz (grey dots). (b) Excitatory *ν*_*e*_(*t*) (top panel) and inhibitory *ν*_*i*_(*t*) (middle panel) population rate predicted by the mean field model when *g*_*gabab*_ increases (see bottom panel, from zero to *g*_*gabab*_ = 0.08nS) for *ν*_*ext*_ = 1.5 Hz. The power spectrum of *ν*_*e*_(*t*) is reported in the inset of the top panel. Dashed line correspond to the stationary mean field model (i.e. with stationary *GABA*_*B*_). The light solid and noisy lines correspond to network simulations with 10,000 neurons.

Fig. 3. We observe that Σ(*ν*_*e*_) is always zero, meaning that the excitatory firing rate *ν*_*e*_(*t*) does not change over time, i.e. the mean field model displays a stable fixed point corresponding to asynchronous network dynamics. On the contrary, for lower external drive *ν*_*ext*_ = 1.5Hz we observe a transition to a non-zero Σ(*ν*_*e*_) for sufficiently large values of *GABA*_*B*_ conductances, indicating that the excitatory firing rate *ν*_*e*_(*t*) changes over time. In Fig. 4b we report the time trace of neural population rates for values of *GABA*_*B*_ conductances in the regime of large Σ(*ν*_*e*_) (i.e. *ν*_*ext*_ = 1.5Hz and *g*_*gabab*_ = 0.08nS). We observe periodic oscillations, at a frequency of *f*_*osc*_ = 4 Hz in the theta range. In the same figure we report the same simulation for the stationary mean field model, that does not account for the dynamics of *GABA*_*B*_ synapses (dashed lines). Importantly, we observe that the stationary mean field (with stationary *GABA*_*B*_ synapses) does not reproduces the emergence of theta oscillations, at variance with the new mean field model. We have verified that the frequency of oscillations depends on the value of the synaptic time scale *τ*_*gabab*_: the longer *τ*_*gabab*_ the slower network oscillations. Notice that the mean excitatory neurons firing rate (over time) is around 1.3 Hz, which is lower than the frequency of oscillations (around 5Hz). This means that neurons do not spike at every oscillation cycle. We have also estimated the number of spikes per neuron during one oscillation by integrating the excitatory neurons firing rate, and we have found that one neuron fires around 0.27 spikes during one cycle of oscillations (i.e. one spike every 4 cycles on average). These results confirm that these oscillations appear in a sparsely firing regime. We then tested the predictions of the new mean field models against direct numerical simulations. We observe that theta oscillations emerge for sufficiently large values of *GABA*_*B*_ also in direct neural network simulations. The amplitude of these oscillations is lower in network simulations with respect to mean field predictions (see light noisy line in Fig.4b) but the frequency of oscillations is almost the same as the one predicted by the mean field model (~ 5 Hz). In Fig. 5 we report the raster plot (Fig. 5a-b) and the power spectrum of population activity *ν*_*e*_ for network simulation with and without *GABA*_*B*_ (Fig. 5c), confirming mean field predictions.

**Fig. 5.**
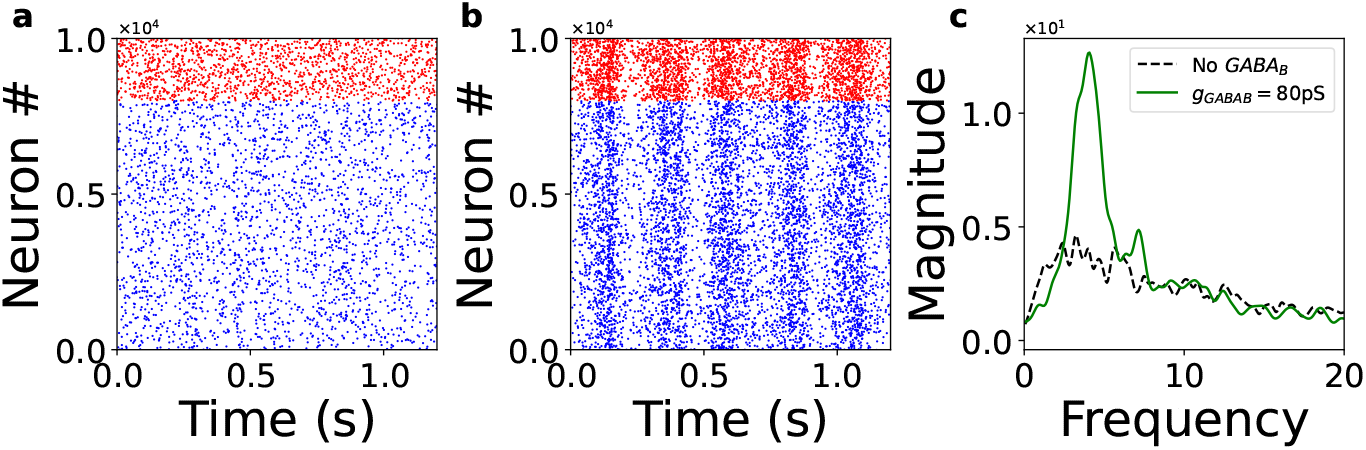
Numerical simulations to confirm mean field predictions of Fig. 4. Panel (a) shows the raster plot obtained from numerical simulation of the neural network with *g*_*gabab*_ = 0pS and panel (b) for *g*_*gabab*_ = 0.08nS and *ν*_*ext*_ = 1.5Hz. Panel (c) shows the power spectrum of the excitatory population firing rate *ν*_*e*_(*t*) for the two conditions.

### 3.5 Distinct roles of increasing inhibition for gamma oscillations

Experimental data suggest that increasing inhibition in cortical circuits can have different impact on population dynamics, and in particular on gamma oscillations. Optogenetically activating interneurons has been found to increase the power of gamma oscillations [28, 29]. On the other side, increasing inhibition by increasing the conductance of *GABA*_*B*_ synapses has seems to have opposite effect, i.e. *GABA*_*B*_ conductance was found to eliminate gamma oscillations in rat hippocampal slices [27]. In this section we show that our mean field model reproduces and predicts this differential role of gamma oscillations. We have thus equipped our mean field model with synaptic delay *τ*_*D*_ = 3ms, while all previous simulations were obtained for instantaneous synapses (*τ*_*D*_ = 0ms). In this version of the model, following the recent work published in [26], we consider an additive noise modeled as an Ornstein–Ulhenbeck (OU) process to Eq.s (8). Accordingly, the external drive *ν*_*ext*_ becomes a time-dependent variable *ν*_*ext*_(*t*) = *ν*_*ext*_ + *ξ*(*t*), where *ξ*(*t*) is an (OU) process evolving according to *dξ* = −*ξθdt* + *σdW*_*t*_ with *θ* = 200Hz, *σ* = 5 and where *W*_*t*_ is the standard Brownian motion. In Fig. 6 we simulate such mean field model with *τ*_*D*_ = 3ms (see the circle in panel a, and the corresponding time trace in the top of panel b). We observe the emergence of gamma oscillations at around *f*_*osc*_ = 80Hz, see top panel in Fig. 6b. Gamma oscillations are not observed in the mean field model for *τ*_*D*_ = 0ms (dashed line, no delay). From this control parameter setup, we have then independently increased the external drive to inhibitory neurons *I*_*ext*_ and the conductance of slow *GABA*_*B*_ synapses in the mean field model. In order to evaluate the amplitude of gamma oscillations we report in Fig. 6a the standard deviation over time of inhibitory firing rate Σ(*ν*_*i*_). Note that this quantity measures the degree of synchrony among interneurons, which contributes to LFP power. The LFP power can indeed be estimated as the sum of net inhibitory and excitatory inputs to pyramidal cells. [36]. In the control condition (circle) we observe a moderate amplitude of gamma oscillations, see the time trace in the top panel of Fig. 6b. By manually activating inhibitory neurons through increasing of *I*_*ext*_ the power of oscillations increases (see the star and the middle panel of Fig. 6b). On the contrary, increasing *GABA*_*B*_ conductance kills gamma oscillations (see the square and the bottom panel of Fig. 6b.

**Fig. 6.**
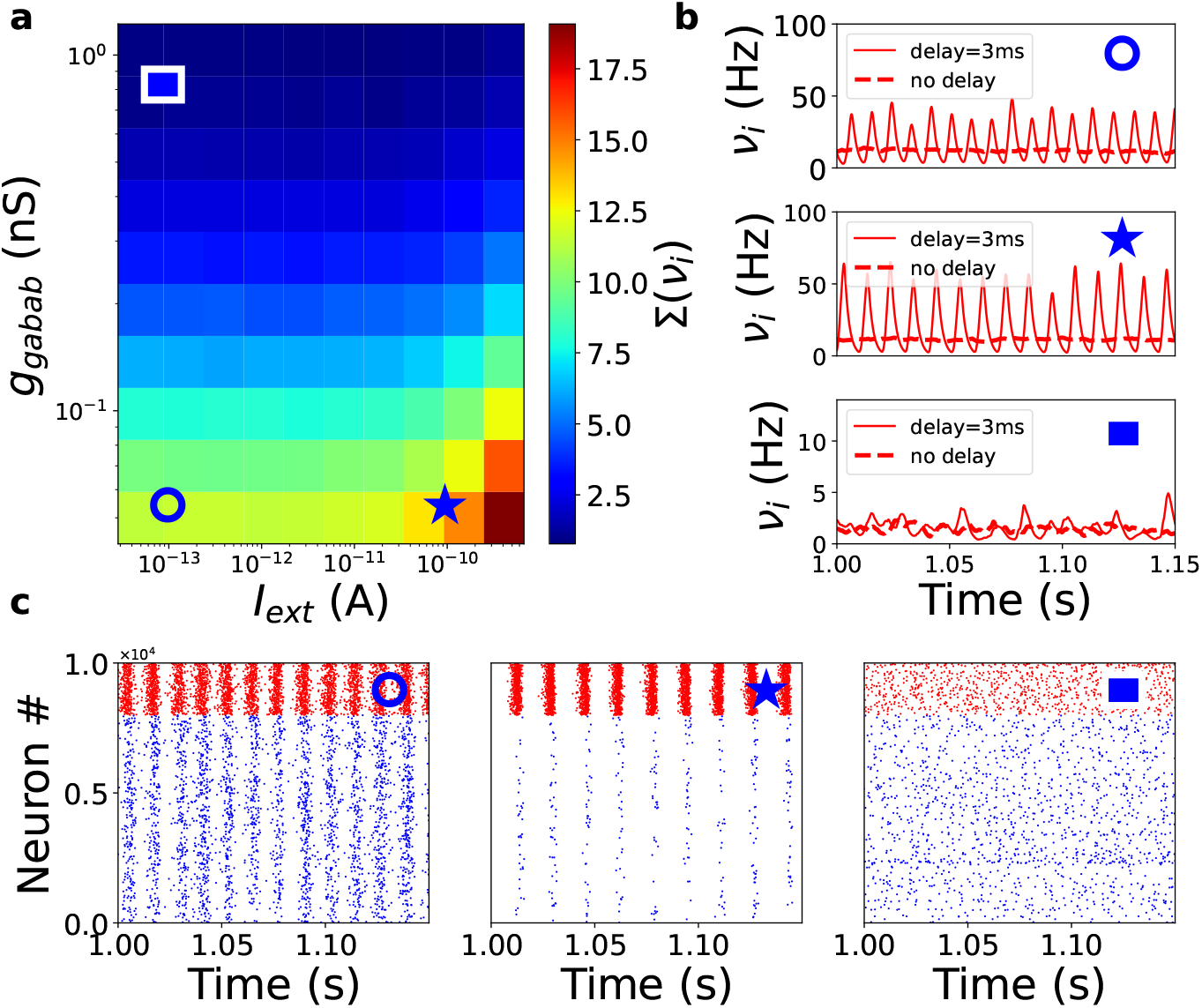
Panel (a) shows the standard deviation over time of the inhibitory firing rate Σ(*ν*_*i*_) for different values of *g*_*gabab*_ and *I*_*drive*_ estimated from simulations of the mean field model with *τ*_*D*_ = 3*ms*. We used *T* = 2*ms* for the mean field simulations. Circle, star and square indicate a set of parameter values employed in panel b (from top to bottom), where we report *ν*_*i*_ over time for the mean field with *τ*_*D*_ = 3ms (continuous line) and *τ*_*D*_ = 0ms (dashed line). In panel c we report results from numerical simulations of the neural network (we report the raster plots) for the parameter values of circle, star and square, from left to right.

We have then verified these predictions in numerical simulation of the neural network. In Fig. 6c we report the raster plot for the parameters in panel b (circle, star and square) of the same figure. We observe that indeed *I*_*ext*_ and *GABA*_*B*_ have a differential role in the generation of gamma oscillations: increase in inhibitory network synchronization for *I*_*ext*_ and asynchronous dynamics by increasing *GABA*_*B*_. These results are in agreement with our mean field model predictions and with experimental data.

### 3.6 Gamma bursts are regulated by GABAB synapses

Oscillations in brain recordings such as LFP are not stationary limit cycles but typically appear as bursts, i.e. short interval of time characterized by few cycles of oscillatory behavior, separated by asynchronous activity. This has been observed for beta [37] but also for gamma oscillations [30, 38]. Oscillations have thus a non-trivial dynamics, characterized by the appearance and the duration of gamma bursts. Our model has shown that *GABA*_*B*_ synapses affect the amplitude of gamma oscillations. In this section we investigate the impact of *GABA*_*B*_ synapses on the appearance of gamma bursts. Also here we consider an additive noise modeled as an Ornstein–Ulhenbeck (OU) process to Eq.s (8), see previous section for details and parameters. In Fig. 7 we report the time-frequency analysis of excitatory neurons’ firing rate predicted by the mean field model. We then superimpose the mean field *GABA*_*B*_ conductance *G*_*b*_. Different panels are obtained for different values of *GABA*_*B*_ synaptic time scale *τ*_*gabab*_. We observe that for sufficiently fast inhibition, gamma oscillations are very powerful and sustained over time, resembling the limit cycles observed in Fig. 6b for low *GABA*_*B*_ conductance, i.e. a situation where inhibition is dominated by fast *GABA*_*A*_ synapses. Increasing the time scale of *τ*_*gabab*_ we observe that gamma bursts become more and more rare and less powerful (lower amplitude). By looking at the trace of *G*_*b*_ we observe that gamma bursts are appearing by the disinhibition from *GABA*_*B*_ release and they terminate after a sufficient level of *GABA*_*B*_ is accumulated by the activation of inhibitory neurons in a gamma burst. We thus conclude that the dynamics of *GABA*_*B*_ receptors and its time scales regulates the appearance and the duration of gamma bursts.

**Fig. 7.**
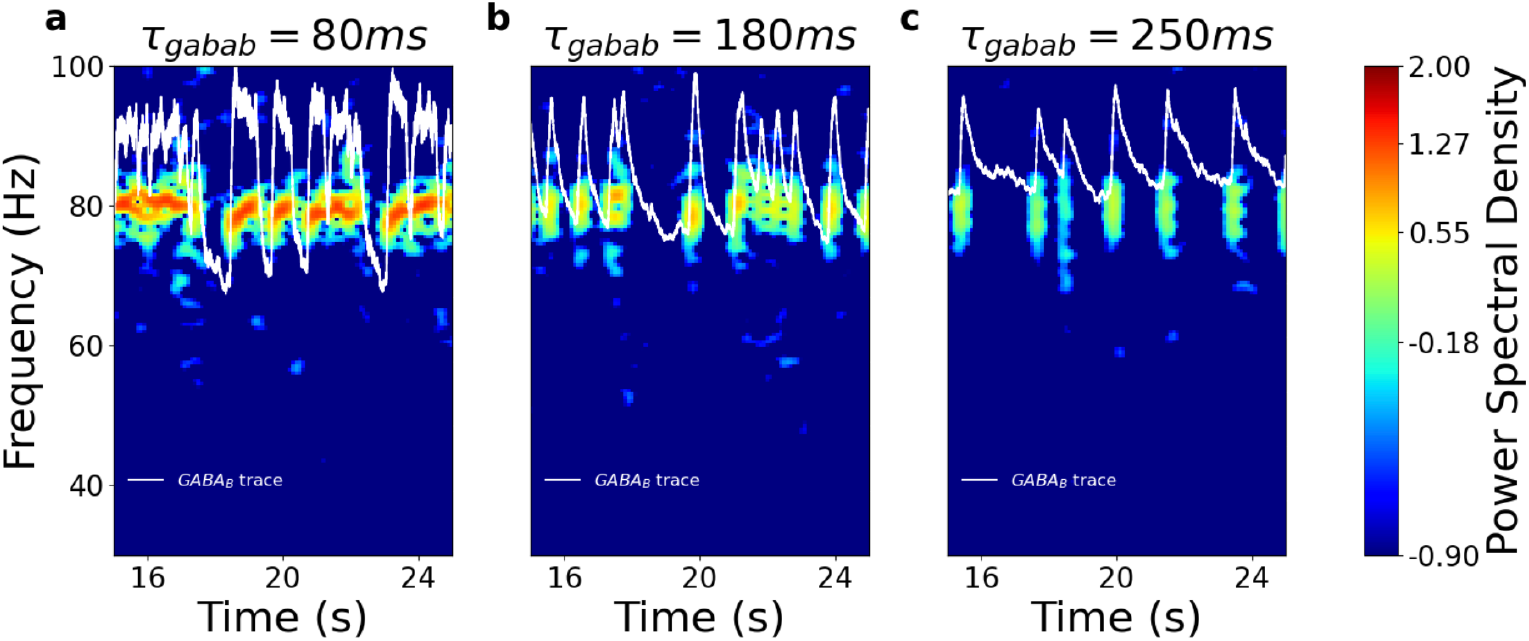
Power spectrograms of the excitatory population rate *ν*_*e*_(*t*) from the mean field model 8 for different values of *GABA*_*B*_ conductance time scale *τ*_*gabab*_ (panels a-b-c). We use the short-time Fourier transform (STFT) subroutine from the signal package of the SciPy library [39] to obtain the Fourier transform of *ν*_*e*_(*t*) within a running time window of length Δ_*t*_ at time t. We perform STFT using a 90% overlap, with a window length of Δ_*t*_ = 0.05s. The spectrogram was visualized using a colormap, where the color code represents the normalized power spectral density. The white trace represents the *GABA*_*B*_ conductance *G*_*b*_. For this simulation *ν*_*ext*_ = 7Hz and *g*_*gabab*_ = 0.01nS.

### 3.7 Nonlinear *GABA*_*B*_ synapses block epileptic seizures

We now employ our excitatory and inhibitory network of neurons and its mean field model to investigate the stability of network dynamics. In particular, we focus on the nonlinearity of *GABA*_*B*_ synapses, i.e. the fact that synapses are activated only if there is a sufficient number of synchronous incoming spikes. The function 𝒢(*ν*_*i*_) follows a sigmoid that increases when *K*_*i*_*ν*_*i*_ *>* 1, that corresponds to more than 10 presynaptic spikes in 100ms. In our previous simulations we considered *K*_*i*_ = 100 and we worked in setups such that the inhibitory neurons firing rate *ν*_*i*_ was usually much higher than 0.01 Hz, thus 𝒢 was always equal to 1. We have indeed verified that the results of previous simulations are unchanged if we consider linear synapses with 𝒢 = 1. Nevertheless, the situation changes if we consider very sparse networks with low activity. In the simulations of Fig. 8 we considered a sparser network with *p* = 0.025%, i.e. mean inhibitory (excitatory) inputs per postsynaptic neuron *K*_*i*_ = 50 (*K*_*e*_ = 200) and a lower external drive *ν*_*baseline*_ = 1.5Hz. This allows to model a sparse network with low activity, where the amount of open *GABA*_*B*_ receptors is low during spontaneous activity. In Fig. 8a-d we have performed a simulation on this network, by including a perturbation at time *t*_*ep*_=3s, where excitatory *AMPA* conductance are suddenly increased (see panel d). Before *t*_*ep*_ = 3s we observe a sparse asynchronous firing of neurons (see panel a and b) and a very low fraction 𝒢 ~ 1% of activated *GABA*_*B*_ synapses (see panel c). In panel a we considered linear synapses, i.e. 𝒢 does not depend on *ν*_*i*_ and is fixed to 𝒢 ~ 1%. We observe that when *g*_*ampa*_ increases at time *t*_*ep*_ there is a transition to high frequency activity (around 150 Hz) characterized by hypersynchronous bursts (see the inset of panel a). This corresponds to a pathological hyperactivity typical of epileptic seizures. In panel b we considered instead the nonlinear *GABA*_*B*_ synapses. We observe that the network does not transit to hyperactivity but to relatively slow oscillations driven by the dynamics of the fraction of activated *GABA*_*B*_ receptors 𝒢 (see blue continuous line in panel c). We have then performed the same simulations in the mean field model observing very similar results. In this case, we considered a lower value of *g*_*ampa*_ for the transition to epilepsy. Data are reported in panels e-g. We observe that linear *GABA*_*B*_ yields hyperactivity at 150Hz (green dashed line in panel e), while nonlinear *GABA*_*B*_ drive the system to healthy slow oscillations driven by the dynamics of 𝒢. Notice that our mean field approach cannot reproduce oscillations as fast as (or even faster) than the time scale of the Markovian process *T* ~ 10ms. As a result, it cannot reproduce the hyper-synchronous oscillations that we observe in the network during epileptic state (see the inset of Fig.8a). In the mean field model, this state is represented by an abnormally high population firing rate *ν*_*e*_ ~ 150Hz. These results show that our mean field model can capture important features of the effect of nonlinear *GABA*_*B*_ synapses, such as their potential role in stabilizing network activity to avoid epileptic seizures.

**Fig. 8.**
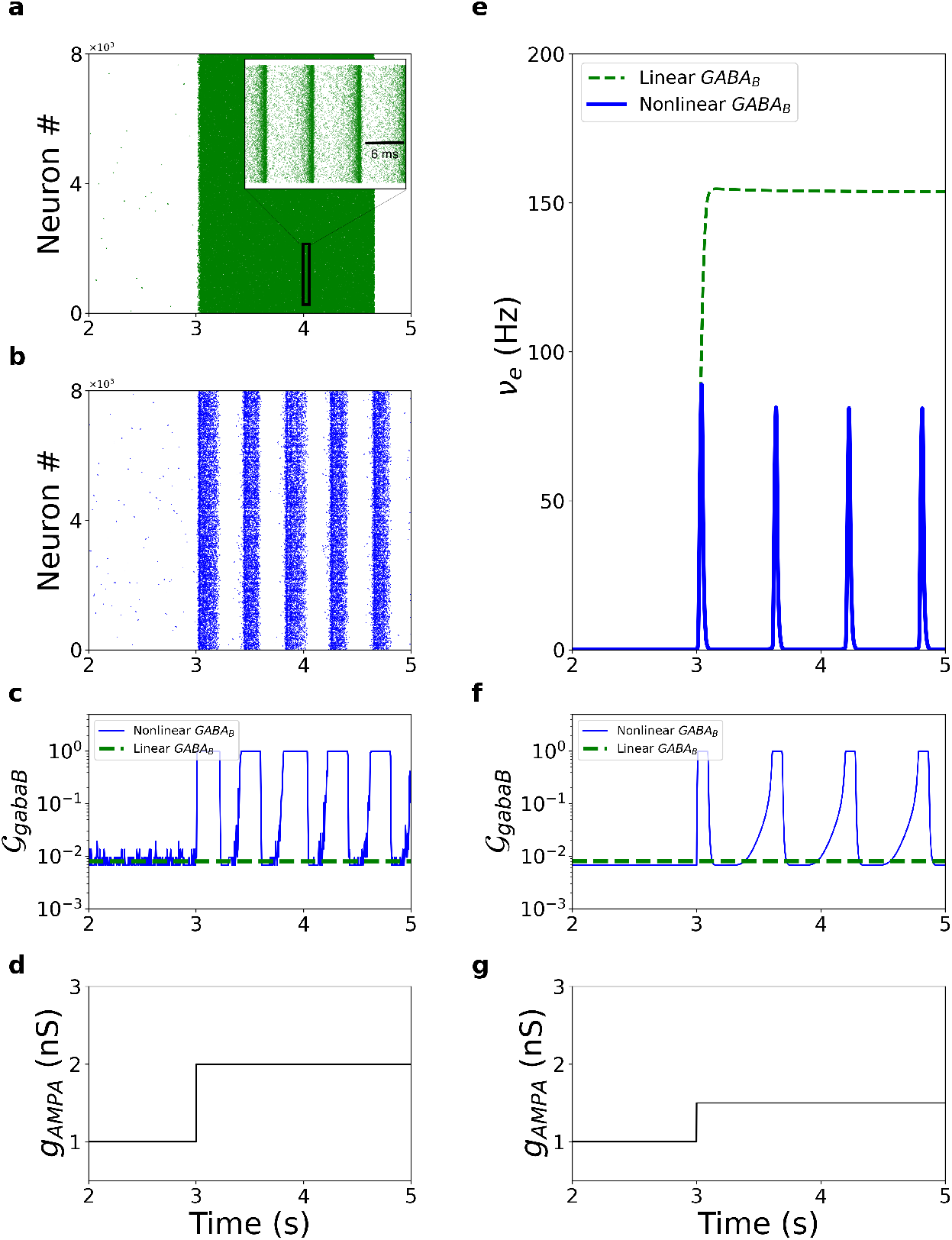
Panels a-d refer to neural network simulations. Panel a shows the raster plot of excitatory neurons for linear *GABA*_*B*_ synapses, i.e. 𝒢 ~ 1%, while panel b shows the raster plot for nonlinear *GABA*_*B*_ synapses, see Methods. In panel c we report the values of 𝒢 for the simulation in panel a (green dashed line) and panel b (blue continuous line). In panel d we show that the values of *g*_*ampa*_, at *t* = *t*_*ep*_ = 3s increases, leading to an epileptic seizure for the linear model (panel a). In the inset of the panel a we report a zoom for a duration of 20ms where we observe hypersynchonous bursts. Panels e-g refer to mean field model simulations. Top panel shows the firing rate of excitatory neurons *ν*_*e*_ for the linear (green dashed line) and nonlinear (blue continuous line) *GABA*_*B*_ conductance model. Panel f shows the values of 𝒢 and panel g the values of *g*_*ampa*_. In these simulations *K*_*i*_ = 50, *ν*_*baseline*_ = 1.5Hz and *b* = 2pA.

## 4 Discussion

In this paper we have developed a new mean field model of sparse conductance-based spiking neural networks including the slow dynamics of *GABA*_*B*_ receptors. The predictions of the mean field models have been directly compared to numerical simulations of a spiking neural network of excitatory and inhibitory neurons. We have found a very good agreement between the mean field model and the networks simulations, both in spontaneous activity and in the response to external stimulation.

A first important outcome of this model is the emergence of theta oscillations at 4-5 Hz. The mean field model predicts that these oscillations can be locally produced by the slow time scale of *GABA*_*B*_ synapses. We have verified that a decrease in *GABA*_*B*_ time scale increased the frequency of these oscillations, covering the theta range observed in rats (4-12)Hz and in Humans (4-8)Hz [40]. The important point of this analysis is that the stationary mean field model (that does not account for the dynamics of slow *GABA*_*B*_ synapses) cannot predict the emergence of these oscillations.

A second important outcome from this mean field model is its capacity to predict different role of increasing inhibition in neural networks. Inhibition can be increased either by directly activating inhibitory neurons (such as in experiments of optogenetics) or by increasing *GABA*_*B*_ conductances (such as by pharmacological agonists). These two manipulations yield different results. Experimental data suggests that *GABA*_*B*_ reduces gamma oscillations, at variance with optogenetic stimulations of interneurons. Our mean field model reproduces gamma oscillations and the different role of these manipulations, thus suggesting that these are nontrivial emergent properties of neuronal networks captured by our “simple” and low dimensional mean field model.

In our approach we have considered *GABA*_*B*_ receptors having a slow decaying activity, at variance with *GABA*_*A*_ synapses. This is a main difference between these two receptor types, but not the only one. In fact, as mentioned in the method section, *GABA*_*B*_ receptors have a nonlinear activation at variance with *GABA*_*A*_ receptors. They get activated only when there are 3-10 repetitive presynaptic inputs in a short time interval of around 100ms [31]. This property could play a crucial role to stabilize the network and avoid epileptic seizures, as our model predicts. Nevertheless, this happens only if we consider very sparse networks with very low firing rates. In our mean field model (and in direct numerical simulations of neural networks) nonlinear *GABA*_*B*_ synapses seem to prevent epileptic seizures by strongly moving to a fully open *GABA*_*B*_ receptor state, characterized by slow oscillations of few Hz driven by the nonlinear opening and closing of *GABA*_*B*_ receptors. As these nonlinear receptors are strongly activated by transient events of strongly active interneurons, it will be interesting in future studies to investigate the impact of bursting interneurons on the emergent dynamics of neural networks through the nonlinearity of *GABA*_*B*_ synapses. In this view, bursting interneurons could serve as openers of *GABA*_*B*_ synapses, putting the network in a safe mode to avoid epilepsy. This could generalize to any type of high firing activity, which would likely activate *GABA*_*B*_ receptors [31].

It has been largely reported that dysfunction of the GABAergic system has a crucial impact on the genesis and the propagation of epileptic seizures [41]. Many studies have shown an impact of *GABA*_*A*_ receptors for the genesis of epileptic seizure in mice models [42]. Our model proposes instead a major role of *GABA*_*B*_ receptors, due to their nonlinearity and slower dynamics. Interestingly, some experimental work points to *GABA*_*B*_ receptors as principal actors for the absence of seizures, presumably in the thalamic nuclei [43, 44]. It is important to notice that the data in this regard are quite complex as they show that *GABA*_*B*_ receptor involvment can be pro or anti epileptsy depending on the nature of the pathological neuronal networks involved [45]. Interestingly, our mechnistic model suggests a major role of *GABA*_*B*_ receptors, but only for sufficiently sparsely connected networks with low firing activity in healthy conditions. Finally, let us point out that on top of GABAergic involvment in epilepsy, many experimental works suggest a major role of glial cells and plasticity [46]. Future computational works should study the distinct or synergetic role of these mechanisms for the stability of network dynamics.

In our model we considered only one type of interneurons, fast spiking inhibitory neurons resembling parvalbumin (PV) cells. Nevertheless, it is known that different interneuons types are present in the cortex [47] and in the hippocampus [48]. Also, recent works seem to suggest that different interneurons types play synergetic role in network dynamics and stability [49]. Future works should aim to develop mean field models able to include the diversity of these interneurons. Our approach, capable to model different neurons proprieties (such as duration of action potential and spike frequency adaptation) seem suitable for this goal. Another important aspect is the ratio between excitatory and inhibitory neurons. We have here considered 80-20% for the sake of generality, but this ratio is variable across brain areas. Future works could employ our model to study the effect of *GABA*_*B*_, for example on seizure susceptibility, on different regions depending on their specific E/I ratio.

Furthermore, the adiabatic approach employed here to model slow inhibitory synapses can be employed also for slow excitatory NMDA receptors. A similar approach has been employed in the context of the exact reduction of QIF neurons [50], by taking also into account the additional nonlinearity of NMDA receptor. This approach could be employed, combined with the one presented here, to include NMDA receptors in the context of sparse conductance based networks such as those employed in this work. This model could be also employed to investigate the effects of NMDA receptors to regulate gamma oscillations [51].

Finally, this mean field approach can be employed to study large scale interaction between brain areas through cross frequency coupling, and the role of receptor specific agonist on interneurons synapses. This approach can show differential role of pharmacological action (namely *GABA*_*A*_ or *GABA*_*B*_ agonists) in the functional connectivity at large scale by using different species of connectomes, following the large scale modeling developed in [10] for the mean field model developed in [25].

## Acknowledgements

We thank B. Delord, E. Procyk and F. Tahvili for useful comments and discussions on this work. This work was supported by the French Ministry of Higher Education (Ministére de l’Enseignement Supérieur) and the project LABEX CORTEX (ANR-11-LABX-0042) of Université Claude Bernard Lyon 1 operated by the ANR.

